# Structure of bacterial communities in Japanese-style bathrooms: Comparative sequencing of bacteria in shower water and showerhead biofilms using a portable nanopore sequencer

**DOI:** 10.1101/2021.07.14.452346

**Authors:** So Fujiyoshi, Yukiko Nishiuchi, Fumito Maruyama

## Abstract

Showers are one of the main exposure routes to diverse microbes for end users in built environments. Bacteria in water are responsible for biofilm formation on surfaces, and the inside of a showerhead is a specific niche. Here, for the purpose of microbial characterization, source estimation and possibility of infection, the bacterial compositions of both shower water and showerhead biofilms in the same bathroom were determined and compared using a portable nanopore sequencer. The results suggest that specific bacteria in source water would primarily adhere to the surface of the showerhead where they subsequently form biofilms, and the community compositions within biofilms largely vary depending on environmental factors. The relative abundance of several pathogenic bacterial genera in both water and biofilm samples was low. We suggest that it is important to manage risk of infection in each household, and rapid on-site analysis of microbial communities will allow the realization.

## Introduction

We are surrounded by numerous microbes in our daily lives, and the bathroom is one of the largest reservoirs of microbes in the built environment due to moderately high temperatures and humidity (Neu *et al*. 2018). The typical Japanese bathing style is to immerse the body in the bathtub and wash the body outside the bathtub, and many Japanese people are known to bath for longer times and with higher frequency (approximately 30 min, every day) than people in other countries (Tochihara, 1999; Odamaki *et al*. 2019).

Shower water is generally supplied by a drinking-water distribution system, so microbes found in the water usually pose no risk for healthy individuals. To control microbial contamination in water, different chemicals and chemical treatment methods (ozone, fluorine, chlorine, chlorine dioxide, monochloramine, and copper-silver ionization and photocatalysis) and physical treatments (thermal inactivation, UV, filtration, and precipitation) are continuously applied (Cervero-Aragó *et al*. 2015; Huang *et al*. 2020). However, some of the microbes in the water can be opportunistic pathogens (Ops) capable of causing serious and life-threatening infections in severely immunocompromised individuals. Some Ops in water cause pneumonia (e.g., non-tuberculous mycobacteria (NTM), *Legionella* spp. and *Mycoplasma* spp.), asthma, or allergies (Dannemiller *et al*. 2016; Montagna *et al*. 2016; Nishiuchi *et al*. 2017). Bacteria in water are important in biofilm formation on surfaces, as they provide the initial cells for attachment and further biofilm development, and the inside of a showerhead is a specific niche that is moist, warm, dark, and frequently replenished with low-level nutrient resources. Bacteria are more concentrated in the biofilm than in the feed water; the number of colony forming units (CFUs) in water after chlorination is 10^2^ CFU/mL, while in biofilms, the number of CFUs is between 10^4^–10^5^ CFU/mL (Kormas *et al*. 2010; Peng *et al*. 2020; Novak *et al*. 2020). In addition, biofilms provide protection against environmental stressors such as antimicrobial agents and disinfectants (Johnson 2008).

Previous microbiological studies of built environment have used culture methodology to detect and identify microbes and have focused primarily on *Legionella pneumophilia* and *Mycobacterium avium* subsp. *hominissuis* (MAH) (Arikawa *et al*. 2019; Falkinham 2020a). Nishiuchi *et al*. reported predominant colonization of *M. avium* in bathtub inlets of bathrooms of patients in Japan using culture isolation (Nishiuchi *et al*. 2007; Nishiuchi *et al*. 2009). Subsequently, Iwamoto *et al*. reported a high degree of genetic relatedness between bacterial isolates from pulmonary MAH patients and bacterial isolates from their bathrooms based on variable numbers of tandem repeats (VNTR) analysis of 19 loci (Iwamoto *et al*. 2012). These reports implied that bathrooms are potentially a major source of MAH in Japan. However, the main limitation of those studies was that it was unclear whether the transmission of the pathogen originated from the patient or the drinking water distribution systems. Thus, understanding the composition of the microbiome in both water and biofilms is of great importance for health risk assessment.

In 2012, a portable nanopore sequencer MinION (Oxford Nanopore Technologies, London, UK) was developed, which is now considered a breakthrough in DNA sequencing technology. The MinION sequencer enables real-time, on-site analyses of any genetic material (Schmidt *et al*. 2016; Parker *et al*. 2017; Nakagawa *et al*. 2019). Compared to the currently widely accepted MiSeq sequencer (Illumina, San Diego, CA, USA), the MinION sequencer can generate a much longer read length, although with lower accuracy. The increased information content inherent from longer read lengths helps researchers with alignment-based taxonomy assignments. Nygaard *et al*. analysed the building dust microbiome using MinION and MiSeq, and showed that MinION had a better taxonomic resolution than MiSeq at the genus and species levels. By optimizing the PCR conditions, MinION was shown to provide accurate microbial community composition and more accurate data than MiSeq in terms of species-level matches (Fujiyoshi *et al*. 2020). Thus, nanopore sequencing is an available and useful tool for understanding the status of public health based on microbial genetic information.

In this study, we used a high-throughput portable nanopore sequencer to determine the bacterial community in shower water and in showerhead biofilms.

## Results

### Sampling and Sequence reads

Fifty samples from 25 residential bathrooms (showerhead biofilm and shower water paired samples, hereafter called biofilm and water samples) were collected from the Hokkaido to Kinki regions in Japan (Figure 1A and 1B). As illustrated in Figure 2, microbes from biofilms were generally clumped and embedded in extracellular material, consistent with biofilm morphology. The DNA yields from the biofilm and water samples were highly variable, and detectable amounts of DNA could not always be extracted. After PCR, 7 biofilm or water samples were not amplified; therefore, 7 pairs of samples were removed for analysis of the pairs. After sequencing, 2 pairs of samples were removed due to a low number of sequences reads. Finally, 32 samples in 16 pairs were analysed in this study. Among the 643,780 raw reads from water samples and 489,734 raw reads from biofilm samples, 343,092 and 228,863 reads, respectively, were analysed after filtration (Supplemental Table S1). The mean operational taxonomic unit (OTU) counts from the water and biofilm samples were 400 and 279, respectively. The number of OTUs was not significantly different between water and biofilm samples (p-value =0.16 > 0.05). Good’s coverage values were greater than 98% for all the samples (Supplemental Table S1).

**Figure 1.**
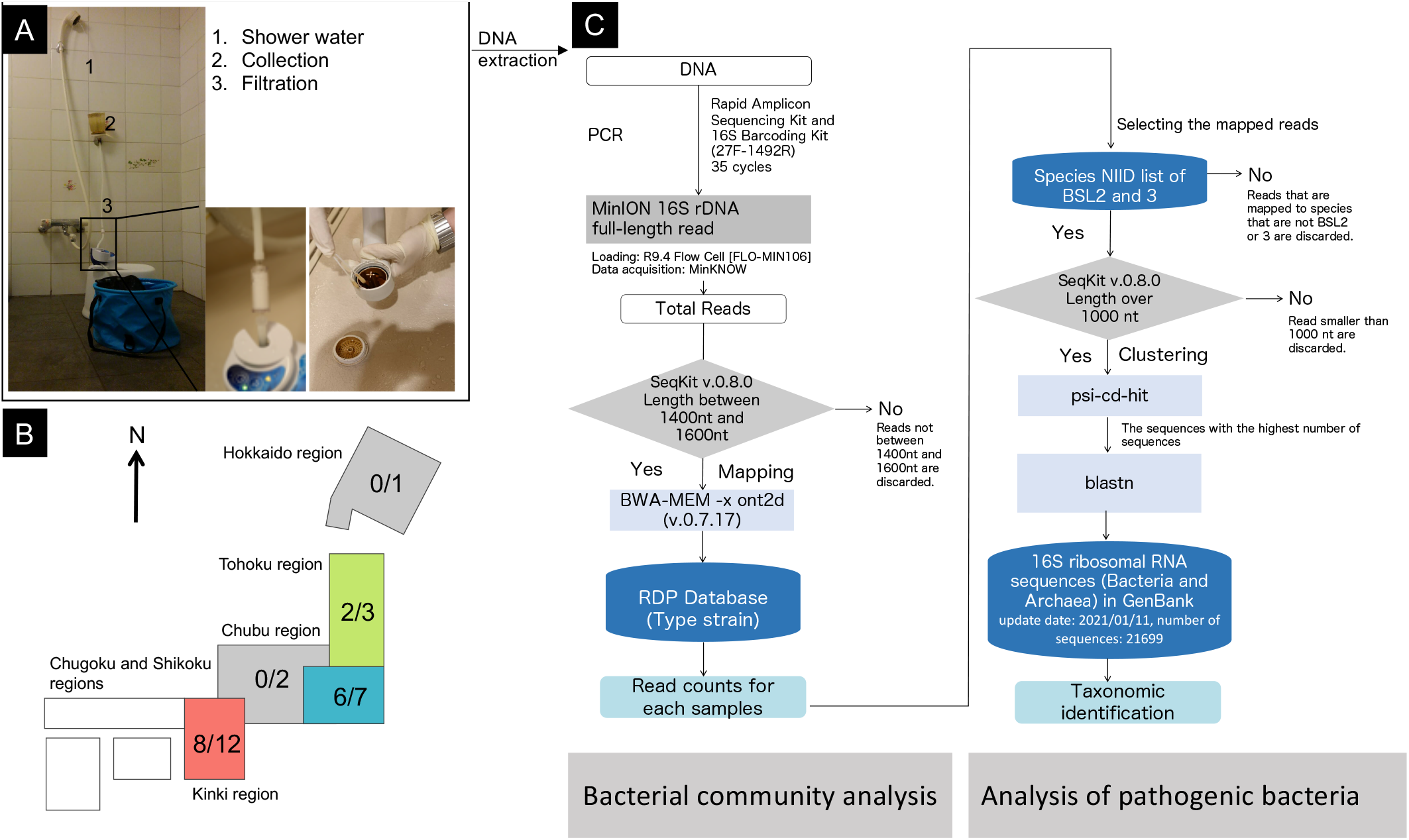
Sampling and sequencing procedure of showerhead biofilms and shower water. Sampling place and method (A). Out of the 25 samples collected from five regions in Japan, 16 samples were analysed. The denominator indicates the number of samples collected, and the numerator indicates the number of samples available for analysis (B). The analysis scheme is divided into two parts: community (left) and pathogen analysis (right) (C).

**Figure 2.**
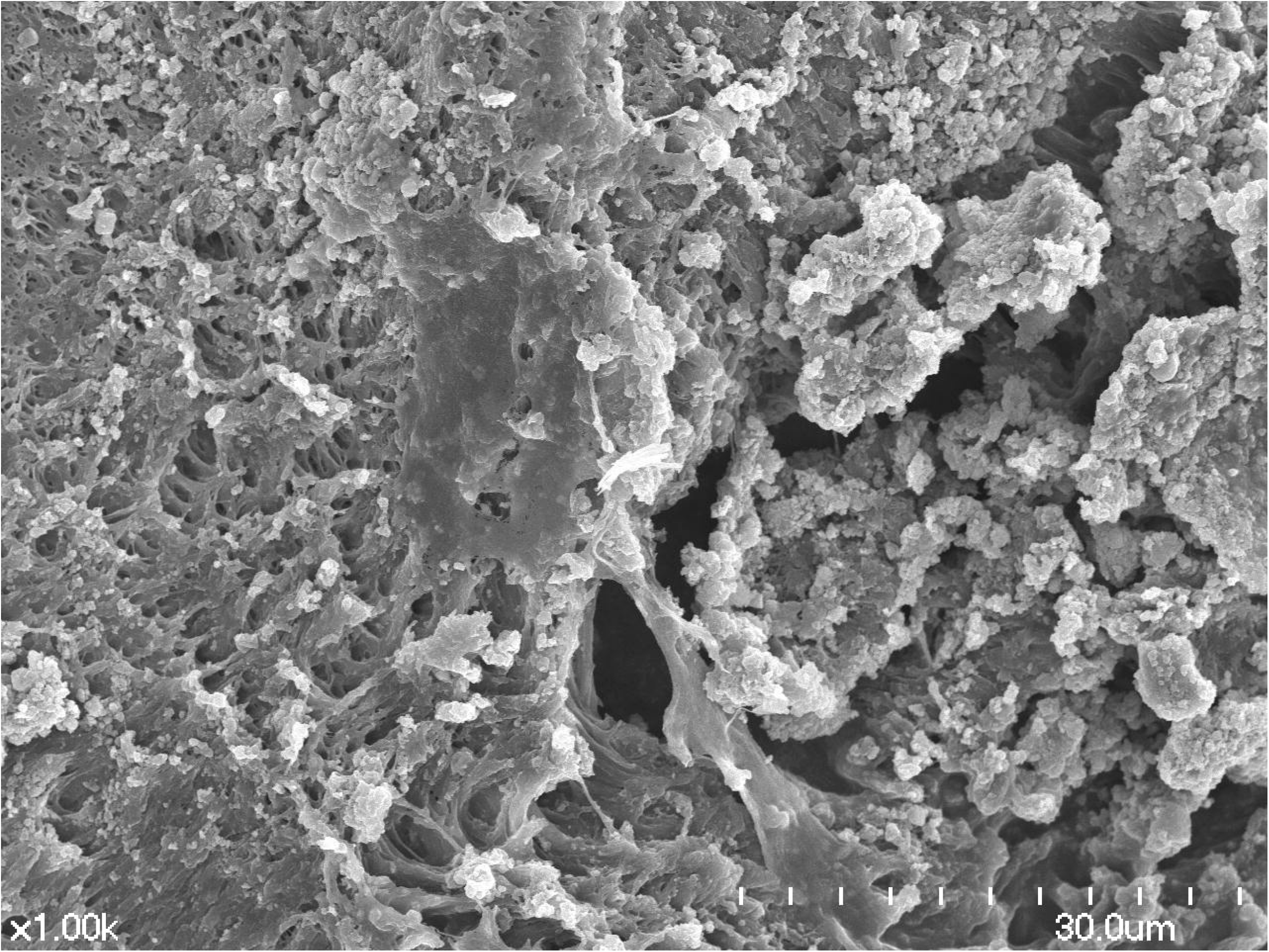
SEM image of a biofilm on the inner surface of a showerhead obtained from one water distributor.

### Comparison of the bacterial community structure in water and biofilm samples

The taxon richness and evenness level reflected by the Shannon index was significantly higher in the water samples than in the biofilm samples (*p* < 0.01) (Figure 3 and Supplemental Table S1). In addition, the Chao1 index, which accounts for species richness, showed that phylogenetic diversity was significantly higher in water samples than in biofilm samples (*p* < 0.01) (Supplemental Table S1). The nMDS plots demonstrated that the bacterial communities in the water samples were more closely clustered than those in the biofilm samples (Figure 4). PERMANOVA results confirmed that the overall community structures were significantly different in both the water and biofilm samples (*p* < 0.01; see Figure 4 for F and R-squared values). Venn diagrams illustrate the percentage of genera shared among each pair of samples. In the biofilm samples, an average of 66.1% of the genera were common to water samples, whereas in the water samples, an average of 36.1% of genera were common to biofilm samples (Figure 5). The percentage of common genera in biofilms was significantly higher than that in water (*p* < 0.01), and the percentage of shared sequences was relatively higher in biofilms (84.1%) than in water (72.1%) (*p* > 0.05).

**Figure 3.**
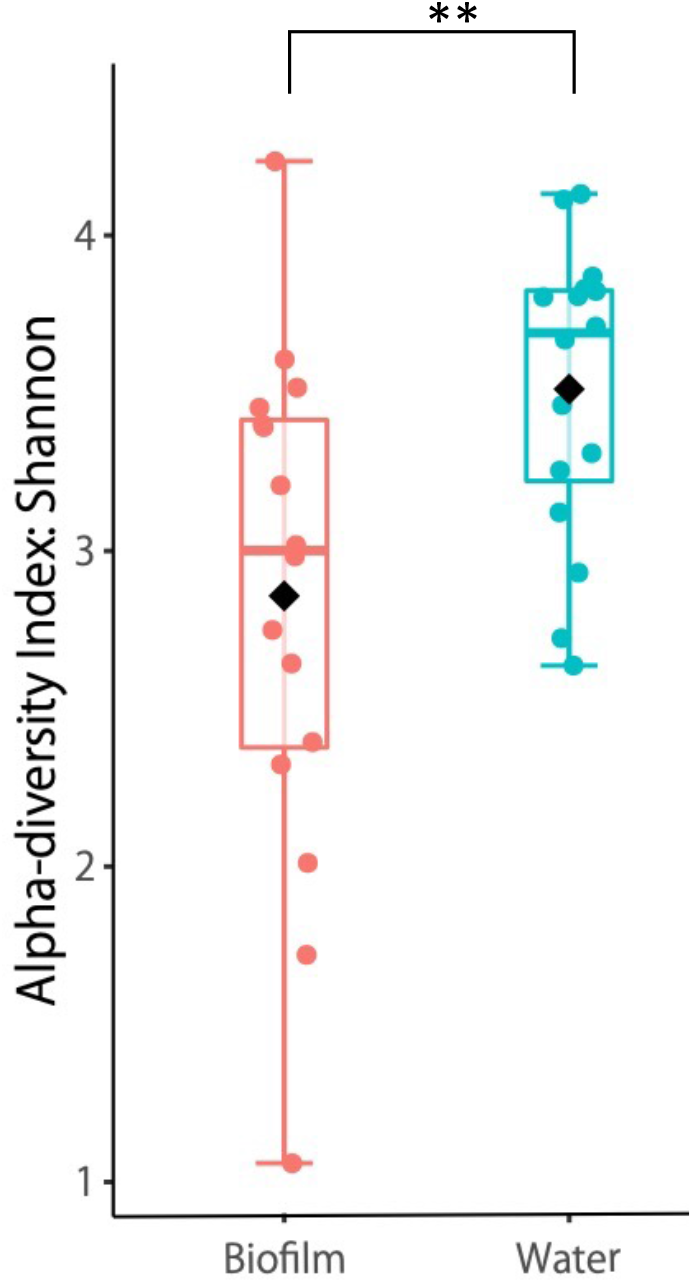
Alpha-diversity measured using the Shannon index. Each box plot represents the diversity distribution of samples from water and biofilm; \*\**p* < 0.01.

**Figure 4.**
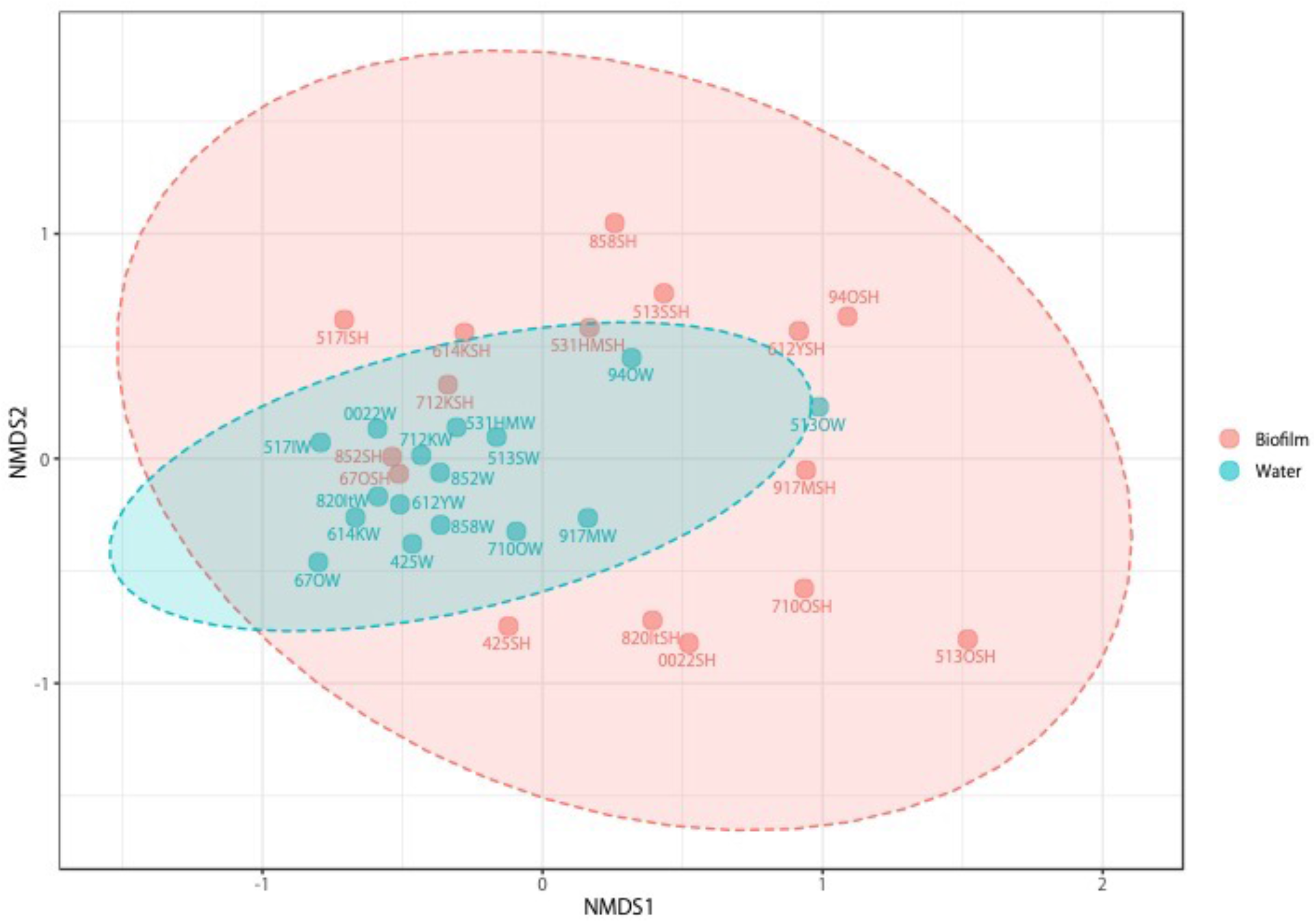
Non-metric multidimensional scaling (nMDS) analysis based on Bray-Curtis dissimilarity metrics of samples labelled according to the sample sources. A clear separation between biofilm bacterial communities and water communities was revealed (PERMANOVA, F-value=3.416, R-squared=0.096, p-value=< 0.01). Ellipses represent the 95% confidence interval. Pink: Biofilm, Blue: water

**Figure 5.**
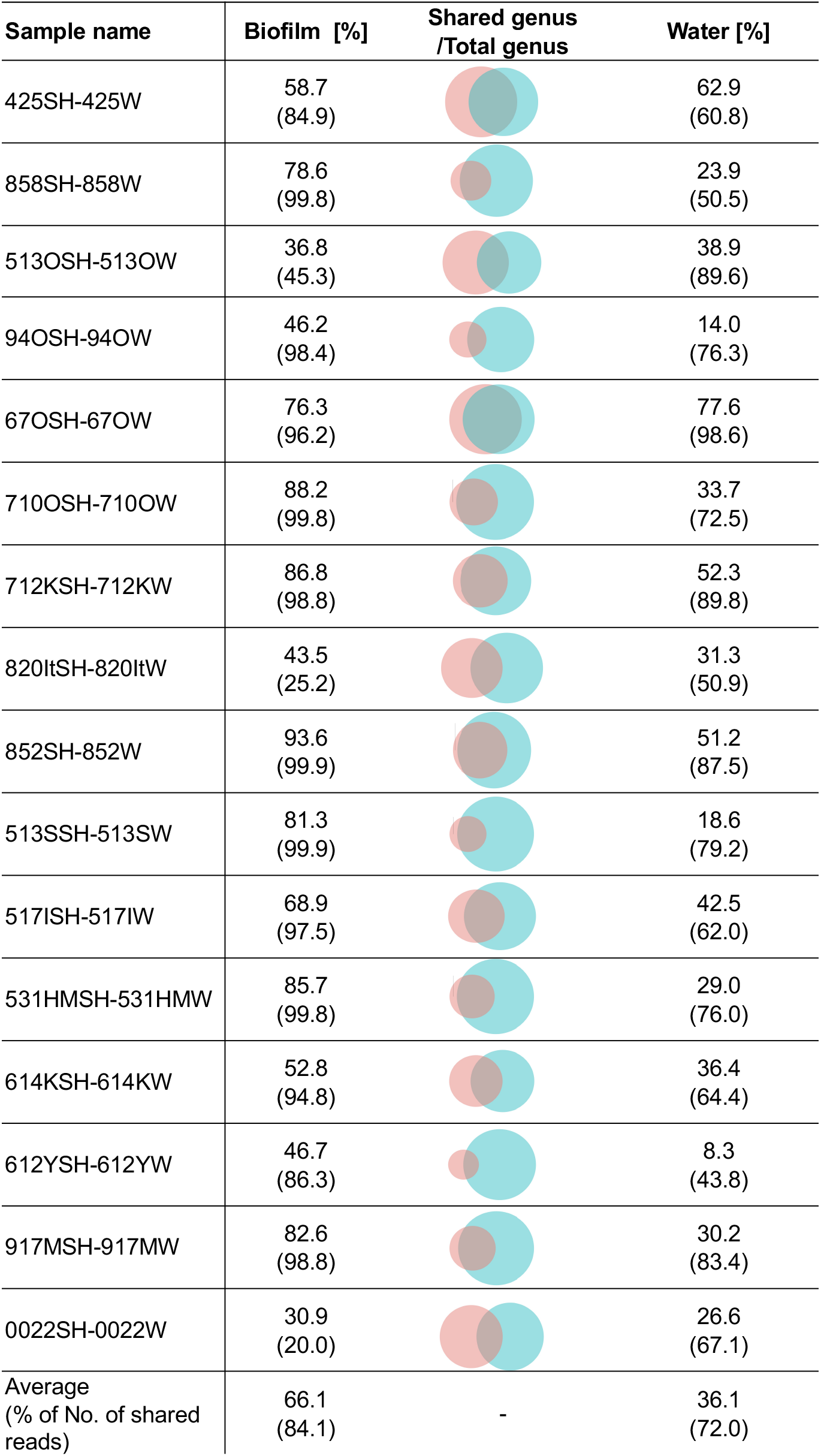
Comparison of the shared genera in biofilm and water samples from the same bathroom. Venn diagrams show a comparison of the percentage of shared genera for biofilms (pink) and water (blue) collected from the same bathroom. Parentheses indicate the percentage of the shared genus sequences in each sample.

### Bacterial community composition

The top three genera in the water samples were *Sphingomonas* (18.8%), *Methylobacterium* (17.0%), and *Phreatobacter* (12.6%), which belong to Proteobacteria (alpha-subclass) (Figure 6), while three genera, *Sphingomonas, Methylobacterium*, and *Bradyrhizobium*, were found in all the water samples. The top three genera in the biofilm samples were *Methylobacterium* (25.2%) and *Sphingomonas* (22.4%), belonging to the phylum Proteobacteria (alpha-subclass), and *Brevibacterium* (8.2%), belonging to the phylum Actinobacteria (Figure 6, for more detail information see Supplemental Figure S1). Only *Methylobacterium* was commonly found in all biofilm samples. To investigate the differences between shower water and biofilm samples, Linear discriminant analysis (LDA) effect size (LEfSe) was performed, and the bacterial taxa significantly associated with each sample was identified. A non-parametric factorial Kruskal-Wallis sum-rank test embedded in LEfSe identified 15 bacterial genera that were characteristic only for water but not for biofilm samples (Figure 7).

**Figure 6.**
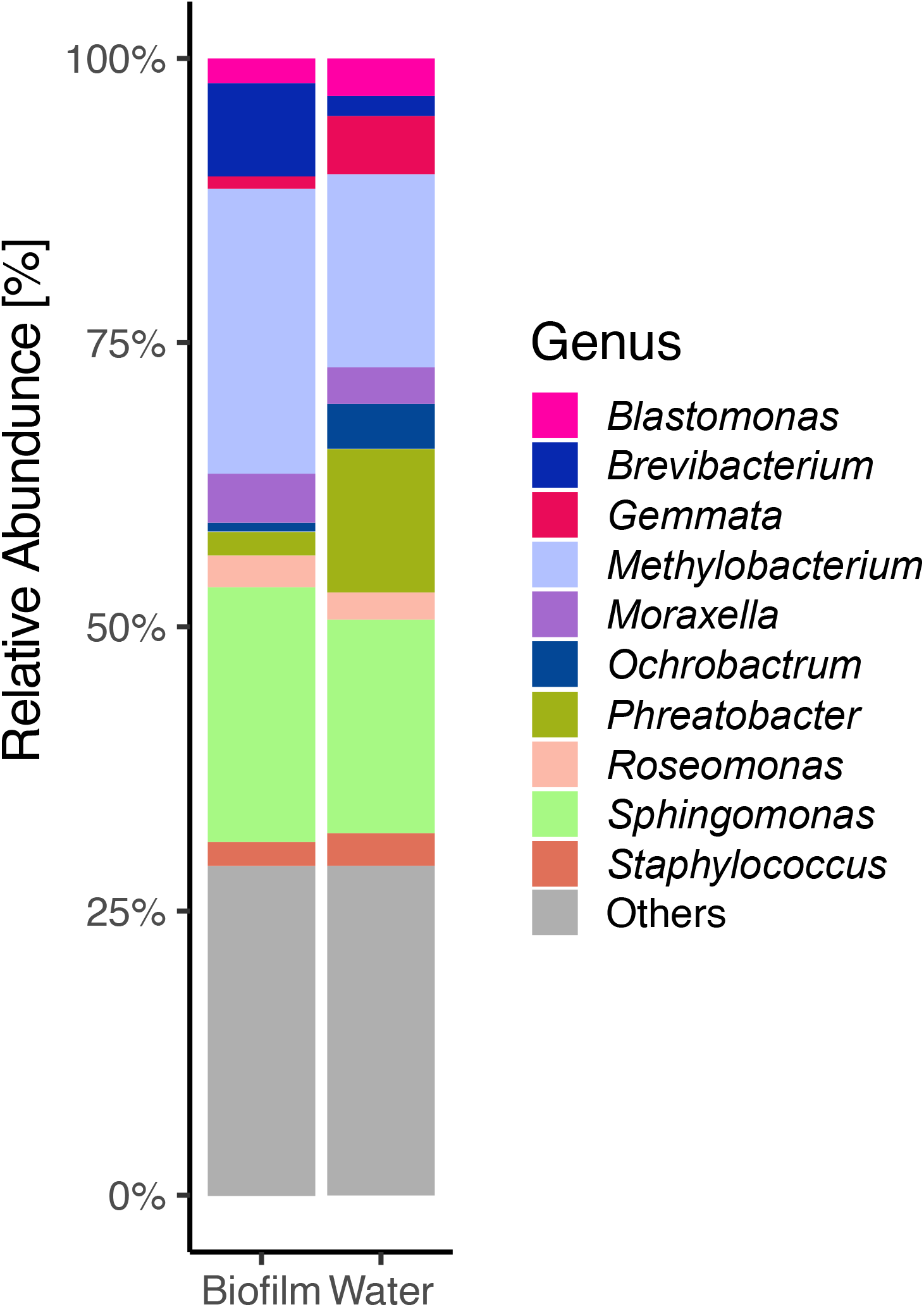
Taxonomic composition of the bacterial community in the biofilm and water samples at the genus level using a stacked bar plot (top 10 genera).

**Figure 7.**
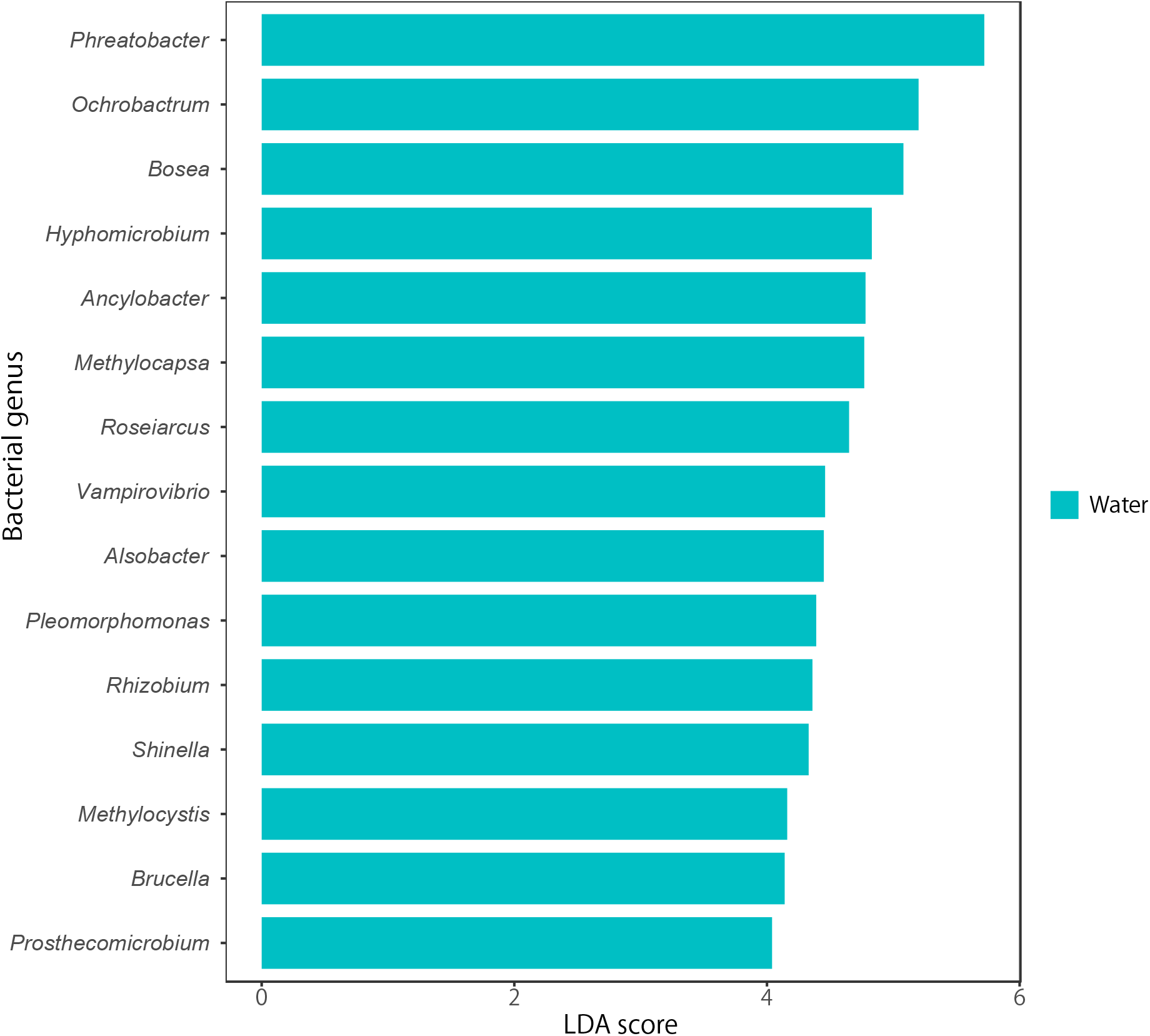
LEfSe analysis of the bacterial community of water and biofilm. Both water and biofilm samples were analysed, but no indicator genera were detected in biofilm. Features are significant based on their adjusted p-value and LDA score. The adjusted p-value cut-off=0.05 and LDA score=4.0.

### Risk of infection in bathroom environments based on the presence of pathogenic bacteria

Based on the BSL2 and BSL3 list from National Institute of Infectious Diseases (NIID), a total of 13 genera were identified in the water and biofilm samples (Figure 8). The genus present at the highest frequency was *Moraxella* with a prevalence of 81% (26/32) and a median value of 0.40%, and the genus with the lowest frequency was *Nocardia* with a prevalence of 3% (1/32) and a median of 0.00%. Only *Moraxella* and *Staphylococcus* had mean values above 1%, and both were detected at high rates in some residences. The median value was 0.00% for most genera. *Mycobacterium*, which includes Ops that cause pulmonary disease, was detected in 34% (11/32) of the samples, with a median of 0.00%. *Legionella* was not detected in this study. Even samples taken from the same bathroom had an Op genus present in the biofilm but not in the water, or vice versa. There was no significant difference in the frequency, mean or median of genera that include pathogens in the biofilm and water samples (p-value > 0.05). At the species level, 5 and 7 BSL2 or BSL3 species were found in biofilm and water samples, respectively. These species accounted for less than 0.30% of the total community (Supplemental Table S2). In addition, these species had a lower percent identity (80.4% average value of identity) when analysed against a more refined and updated BLASTn database (Supplemental Table S3). The species that appeared in both analyses were *Staphylococcus aureus* (87.98% maximum value of identity) and *Vibrio parahaemolyticus* (87.86% maximum value of identity). The genera that appeared in both analyses were *Corynebacterium* (86.26% maximum value of identity) and *Mycobacterium* (86.87% maximum value of identity). For the four species of *Brucella* found in this study, most of them were assigned to different genera with low percent identity (76.34% average value of identity).

**Figure 8.**
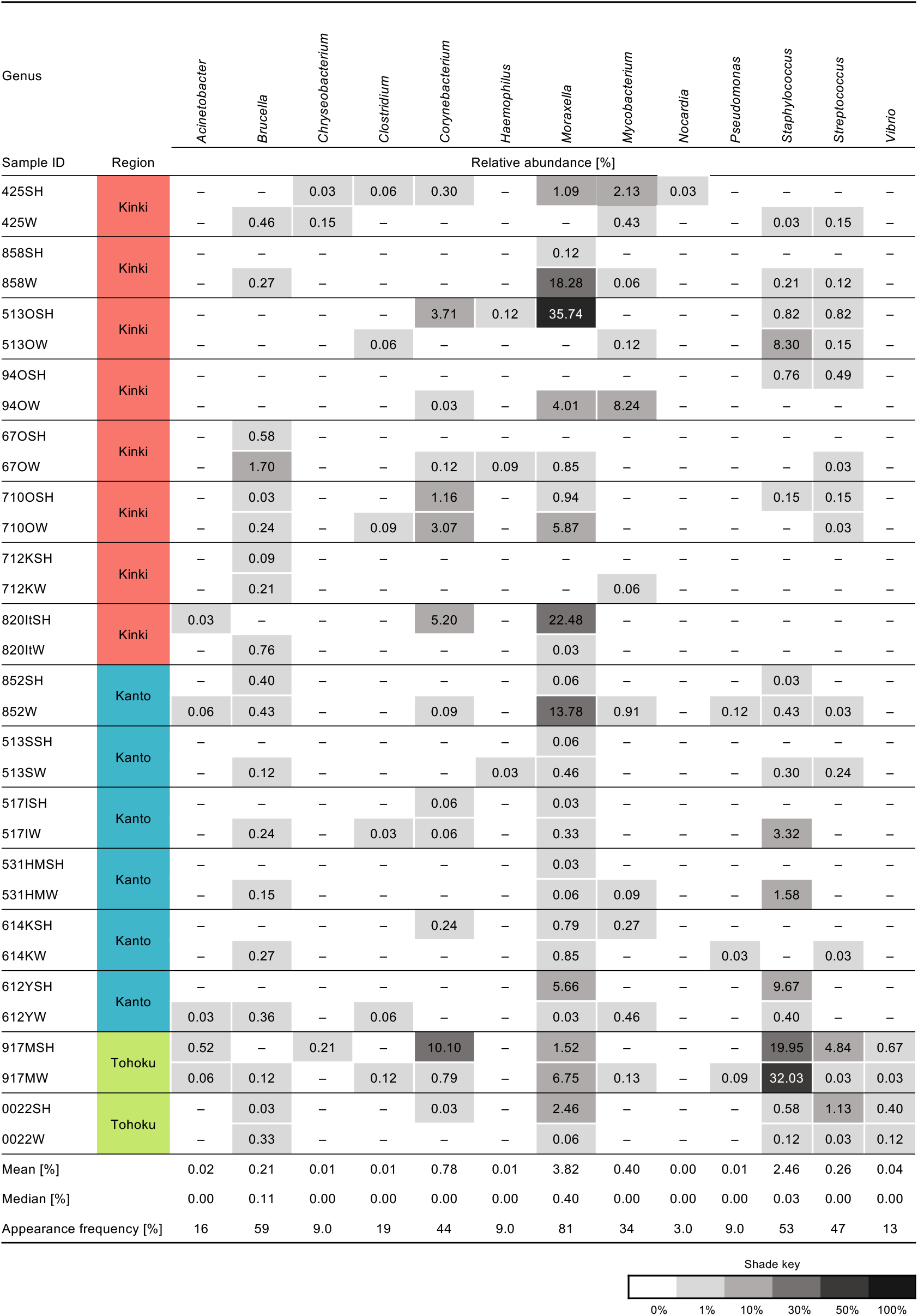
This heatmap table summarizes the results for all showerhead swab and shower water libraries pooled at the genus level and grouped by region in Japan. Figure footnotes: – signifies that the genera were not detected in the sample; SH indicates a biofilm sample and W indicates a water sample.

## Discussion

### Biofilm bacteria are derived from the water

The biofilm formed on the inner surface of the shower head interacts with the shower water (Rasmus *et al*. 2002; Peng *et al*. 2020). The results of our Venn diagram also show that some genera are shared between biofilm and water samples (Figure 5). However, the bacterial community structures of biofilm and water samples were significantly different, even when the samples obtained from the same bathroom (Figures 3 and 4). The bacterial community in biofilms had a lower alpha-diversity than the community in the water samples, suggesting that some, but not all of the bacteria in the water probably adhere to the showerhead surface and subsequently form a biofilm. Beta diversity analysis revealed significant differences between water and biofilm samples (Figure 4). The bacterial communities in the water were more closely clustered than those in the biofilm samples. The quality of drinking water in Japan is standardized by the Ministry of Health, Labour and Welfare, based on a total of 51 items, such as general bacterial CFU, chlorine and cadmium concentration, and each standard has a maximum value below which health is not affected (Nishiuchi *et al*. 2017; Novak *et al*. 2020). All the water samples collected in this study were supplied from drinking water distribution systems (not well or river water), and the well-controlled water quality in Japan may affect the similarity of the bacterial communities in the water samples. However, the bacterial communities in the biofilms differed greatly among samples. This probably suggests that the bacterial community in a biofilm is influenced by factors other than the water, such as the building type, the method of use, and the residents (Ji *et al*. 2017; Proctor *et al*. 2018). In this study, there was no correlation between the material of the showerhead and the bacterial community structure (data not shown).

The bacterial communities examined in this study are consistent with other studies showing the dominance of similar genera in freshwater systems and biofilms in bathrooms (Figure 6) (Feazel *et al*. 2009). *Methylobacterium* spp. are famous for forming pink-coloured biofilms and are prevalent in domestic water-associated environments, such as drinking water distribution systems, shower curtains, and showerheads worldwide (Gallego *et al*. 2005; Zhou *et al*. 2021). In Japan, *Methylobacterium* are also common bacteria with known culture methods (Yano *et al*. 2013; Kawai *et al*. 2019). *Sphingomonas*, generally the most abundant genus detected in the biofilm samples, are ubiquitous in the environment, such as in soil, water, and sediments, as well as on shower curtains and are known to be constantly and consistently present in biofilms even after chlorination (White *et al*. 1996; Kelley *et al*. 2004; Douterelo *et al*., 2018). *Phreatobacter* has been identified as a dominant genus in drinking water distribution systems in China (Li *et al* 2020; Ma *et al* 2020; Jing *et al*., 2021). *Brevibacterium* from biofilms are not as commonly detected in bathrooms as the other three genera. *Brevibacterium* species are isolated from various habitats, such as fermented food, animal and human skin, insects, soil, and mural paintings. Some species, such as *B. casei* and *B. epidermidis*, encountered in human skin, wounds, and blood can cause rare human infections, making them Ops (Denis and Irlinger, 2008). It should be investigated in the future whether the high percentage of *Brevibacterium* in showerhead biofilms is unique to Japan or due to environmental selective pressure. *Sphingomonas*, *Methylobacterium*, and *Bradyrhizobium* were found in all the water samples, while only *Methylobacterium* was found in all the biofilm samples. This shows that the bacteria in the water samples were relatively consistent, but the biofilm constituent bacteria varied among samples. LEfSe analysis identified significant differences in taxa between biofilm and water samples. A comparison of sample sources indicated that there were 15 taxa only from the water samples with LDA scores greater than 4.0 (Figure 7). No taxa identified were considered indicators of biofilms. This suggests that the bacteria in the biofilm samples are of water origin. A total of 66.1% and 36.1% of genera were common between the biofilm and water samples, respectively (Figure 5). These results suggest that specific bacteria in the original water would primarily adhere to the surface of the showerhead and subsequently form biofilms.

### Potentially pathogenic bacteria were present in shower water and biofilm samples, but only a low percentage of them were present in Japanese bathrooms

The Ops found in the bathroom samples are well reported (Kelley *et al*. 2013; Novak *et al*. 2020; Falkinham. 2020b). *Acinetobacter*, *Stenotrophomonas, Pseudomonas, Mycobacterium* and *Legionella* are commonly presented Ops worldwide. In this study, we encountered two pathogen-like species, *Staphylococcus aureus* (87.98% maximum value of identity) and *Vibrio parahaemolyticus* (87.86% maximum value of identity). *S. aureus* is known as a commensal bacterium and as a pathogen that causes opportunistic infections mainly of soft tissues, skin, and wounds (Kozajda *et al*. 2019). Studies have shown that the detection rate is higher in areas with high hand and foot contact (Ojima *et al*. 2002). Infection with *V. parahaemolyticus* can cause gastroenteritis, septicaemia, and infection. It is a seafood-associated and a water-borne food-mediated bacterium (Liu *et al*. 2015). This bacterium has been detected in freshwater in some cases (Maje *et al*. 2020; Silva *et al*. 2018). In addition to the presence of these bacteria, we also detected some Op genera that are closely related to *Corynebacterium* and *Mycobacterium*. *Corynebacterium*, a known human-associated bacterium, is commonly found in bathrooms, according to meta-analysis data from high-throughput amplicon analysis as well as from culture methods (Allen *et al*. 2004; Adams *et al*. 2015). In this study, *Legionella*, a problem in water systems around the world, was not detected. Some studies show that planktonic and biofilm *Legionella* concentrations are reduced by the presence of residual disinfectants within water material (Waak *et al*. 2018; Fish *et al*. 2020). The standard value of residual chlorine in water in Japan is set at 1 mg/L or less, is often 0.1 mg/L or less and is strictly controlled. The fact that *Legionella* was not detected in this study may indicate the high-performance level of the water quality management system in Japan. *Mycobacterium* was detected in 34% (11/32) of the samples, and the mean and median values were 0.40% and 0.00%, respectively (Figure 8). Arikawa *et al*. surveyed *Mycobacterium avium* subsp. *hominissuis* (MAH) in Japan, and the detection rate of MAH was 16.1% (Arikawa *et al*. 2019). Using a culture-independent method, quantitative PCR, Ichijo, *et al*. detected the frequency of *Mycobacterium* spp. in showerheads. The detection rate was lower than that in our study (13%) (Ichijo *et al*. 2014). According to Gebert *et al*., the detection rate of *Mycobacterium* spp. is 37% across households in the United States and Europe, which is similar to the results of this study. However, the relative abundances of mycobacteria were much higher than those in this study, at approximately 15% and 7%, respectively (Gebert *et al*. 2018). Additionally, Feazel *et al*. reported that *M. avium* was identified in 20% of showerhead swabs in the US (Feazel *et al*. 2009). *Mycobacterium* was not present at a higher rate in Japan than in the US and Europe; however, the incidence rate of nontuberculous mycobacterial pulmonary disease in Japan was reported to be 14.7 cases per 100,000 people in 2014 (Namkoong *et al*. 2016), and the number of infected people is increasing rapidly compared to other countries (Yano *et al*. 2017). Non-tuberculous mycobacteria include nearly 200 species that can differ with respect to their ecology and pathogenicity (Tortoli 2014; Falkinham 2020a). Thus, to obtain more detailed information on specific mycobacterial species, it is necessary to sequence the gene using mycobacterium-specific primers, such as *hsp65* (65-kDa heat shock protein), as well as performing identification through culture methods (Telenti *et al*. 1993). In this study, we encountered the assignation of *Brucella* to other genera. Misidentification of *Brucella* species has been reported in many studies (Elsaghir and James 2003; Horvat *et al*. 2011; Carrington *et al*. 2012), except for in those using near-full-length 1,412-bp nucleotide sequences of 16S RNA genes and Sanger sequencing (Gee *et al*. 2004). Because of these recurring misidentifications, an increasing number of laboratories are now relying on molecular methods such as Gram staining, semisolid-medium motility tests and flagellar staining to identify *Brucella* (Yang *et al*.2013). In the results of the sequence analysis in this study, even if *Brucella* was detected, it was necessary to validate the results by molecular methods using cultured strains.

One of the major advantages of using targeted metagenomic techniques, such as 16S rRNA gene sequencing, is that these techniques are culture-independent and can theoretically recover almost all bacterial taxa in any habitat. Therefore, it is possible to identify the microorganisms, including the percentage of potential pathogens, in the entire community. Moreover, MinION can generate long read length. The increased information content inherent from longer read lengths assists researchers with alignment-based taxonomy assignment (Wommack *et al*. 2008). Mitsuhashi *et al*. and Nakagawa *et al*. reported that 5-min and 3-min running times on MinION, respectively, were sufficient to detect specific bacteria (Mitsuhashi *et al*., 2017; Nakagawa *et al*.,2019). Furthermore, by optimizing the PCR conditions, MinION was shown to provide accurate microbial community composition and more accurate data than Miseq in terms of species-level matches (Fujiyoshi *et al*. 2020). However, because of the widespread use of DNA sequencing technology, it is necessary to carefully investigate the analysis methods, databases, and results of these methods before applying them. The tendency is to use top hits to classify microorganisms, but in general, cut-offs of 95% and 98.7% are used to classify bacterial isolates at the genus and species levels, respectively (Stackebrandt and Goebel 1994). This point needs to be discussed carefully, especially when mentioning Op bacteria in samples. Additionally, DNA does not provide information whether a microorganism is alive, dead, or infectious. Therefore, it is necessary and efficient to conduct epidemiological studies using both metagenomic approaches and culture methods simultaneously. In 2020, we published a paper on a “suitcase lab” in which a single suitcase can contain all the necessary equipment from sampling to detection by the LAMP method and showed that the work can be completed on-site within two hours if the specific target species is decided upon (Fujiyoshi *et al*. 2021). In the future, if such a tool can be used to analyse the microbiome on-site quickly and easily, it will contribute to the improvement of microbial risk assessment and control not only of water quality but also of the built environment.

Our comprehensive analysis of the bacterial community in the built environment revealed that specific bacteria in source water would primarily adhere to the surface of the showerhead where they subsequently form biofilms. Moreover, the bacterial communities in biofilms differed greatly among samples. This probably suggests that the bacterial community in a biofilm is influenced by factors other than the water. The findings of this study are the first data in Japan for assessing microbial communities in built environments. This is also the first study to analyse the microbial communities of both water and its biofilm obtained from same bathrooms, by which the microbial similarity and difference of water and biofilm were emphasized clearly. We suggest that it is important to manage risks of infection in each household and that this requires rapid and easy on-site identification of microbial communities.

## Methods

### Sampling

A total of 50 showerhead biofilms and shower water samples from 25 independent bathrooms were collected from five regions in Japan: 3 from Tohoku, 2 from Chubu, 7 from Kanto and 12 from Kinki, as shown in Figure 1B and Supplemental Table S1 (the sampling sites were classified according to Arikawa *et al*. 2019). The showerhead biofilm and shower water samples in the same bathroom were aseptically collected. Showerhead biofilms were swabbed with sterilized nylon swabs (FLOQ swab 552C; Becton, Dickinson and Company, Tokyo, Japan), and 2 L of shower water was filtered on-site using a portable peristaltic pump (Sentino microbiology pump, Pall Life Science, MI) and a 0.2-μm filter cartridge (Sterivex, Millipore, MA, USA) (Figure 1). The cartridge was put into a sterile tube, and then the samples were immediately placed in a cool box at 4°C, transported to the lab within a few hours and kept at −20°C until use.

### Scanning electron microscope (SEM)

The biofilms on coverslips and filters were prefixed in a solution of 2.5% glutaraldehyde in 0.1 M phosphate buffer (PB; pH 7.4) for 10 min and rinsed three times with PB. Samples were then fixed again with 2.5% glutaraldehyde for 1 h and rinsed three times with PB. Another fixation reagent, 1% (w/v) osmium tetroxide in PB, was added to the samples, followed by incubation for 1 h. Subsequently, the samples were rinsed three times with PB and dehydrated with increasing concentrations of ethanol (30%, 50%, 70%, 90%, 99%, and 100%). Dehydrated samples were soaked in isoamyl acetate, successively critical-point-dried with an HCP-2 (Hitachi Ltd., Tokyo, Japan), and coated with 8:2 platinum–palladium alloy using an E-1030 ion sputter (Hitachi Ltd., Tokyo, Japan). The resultant coating was 12 nm thick. Samples were observed using an S4700 scanning electron microscope (Hitachi Ltd.).

### DNA extraction and PCR conditions

All samples were recovered using aseptic techniques and appropriate negative and positive controls. A swab and a filter, picked out from the cartridge, were directly placed in a bead tube of a DNeasy PowerBiofilm Kit (QIAGEN, Germantown, MD, USA) under a laminar flow cabinet, and DNA was extracted according to the manufacturer’s protocol with some modifications (Arai *et al*. 2018): instead of glass beads in a PowerBiofilm bead tube, 400 μL of sterilized φ0.5 mm zirconia beads (TORAY, Tokyo, Japan) and two gains of φ5 mm zirconia beads (TORAY) were used for homogenization. The samples were bead beaten with a bead crusher (TissueLyser II, QIAGEN) at 3,200 rpm for 10 min. The DNA was eluted in 100 μL of elution buffer and then purified and condensed with a Dr. GenTLE precipitation carrier (Takara BIO, Tokyo, Japan). The concentration and purity of the DNA were measured with a DS-11FX+ Spectro/Fluorometer (DeNovix, Wilmington, USA) and a QuantiFluor™ dsDNA System (Promega, Madison, USA). PCR amplification and barcoding of 16S rRNA genes were conducted using the 16S Barcoding Kit (SQK-RAB204; Oxford Nanopore Technologies, Oxford, UK) containing the 27F/1492R primer set and MightyAmp DNA polymerase Ver.3 (Takara Bio). PCR was performed according to a previous report (Fujiyoshi *et al*. 2020). A reaction containing no template served as the negative control. The amplified fragments were separated in a 2% agarose gel, stained with Safelook Load Green (Wako Chemicals Co. Ltd, Osaka, Japan), and checked with a FAS Nano Gel Document System (NIPPON Genetics, Tokyo, Japan).

### Nanopore sequencing library construction

After purifying a PCR product (50 μl) with 30 μl of Agencourt AMPure XP (Bechman Coulter, Tokyo, Japan), the amount and purity of the DNA eluted with 10 μl of buffer solution (pH 8.0, 10 mM Tris-HCl with 50 mM NaCl) were determined as above. One hundred or 50 fmol of purified amplicon DNA was used as input DNA for MinION-compatible libraries. The amplicons were added to 1 μl of rapid adapter (Oxford Nanopore Technologies) and incubated at room temperature for 5 min.

### Sequencing and data analysis

Each nanopore sequencing library was run on FLO-MIN106 R9.4 flow cells (Oxford Nanopore Technologies) after performing platform QC analysis. The amplicon library (11 μl) was diluted with running buffer (35 μl), with 3.5 μl of nuclease-free water, and with 25.5 μl of loading beads. A 48-h sequencing protocol was initiated using MinION control software MinKNOW v.1.11.5 or 1.14.1. MinION sequence reads (i.e., FAST5 data) were converted into FASTQ files using Albacore v.2.1.3 or 2.3.3 software (Oxford Nanopore Technologies). FASTQ files were analysed as described previously (Fujiyoshi *et al*. 2020). The files examined the sequence read length distribution using FastQC (v 0.11. 2) (Andrews. 2010), and Seqkit 0.8.0 (https://bioinf.shenwei.me/seqkit/) (Shen *et al*. 2016) was used to filter the sequence data by the lengths of 1,400–1,600 to include 1,500–base reads. After filtering, the sequence reads were mapped using bwa-mem (v. 0.7. 17) (Li 2013), with the MinION analysis option (-x ont2d) (Jain *et al*. 2015), to a database derived from the Ribosomal Database Project (RDP Release 11, Update 5, Seqt. 30. 2016) (Cole *et al*. 2014), and the top hit was used for genus and species assignment. The RDP hierarchy browser (http://rdp.cme.msu.edu/hierarchy/hb_intro.jsp) was used with the following filters: strain = “Type”; source = “isolates”; size “>= 1,200”; quality = “Good”; and taxonomy = “Nomenclatural” (for analytical scheme, see Figure 1C). Sequences from all the samples were normalized to the sample containing the lowest number of reads (5115 reads). After the removal of singleton OTUs, all data analysis was carried out with R (v. 3.3.1) (R Core Team 2018). The R package vegan (v. 2.5-5) was applied for diversity and community analyses. Two metrics of alpha diversity were used in this study, Shannon diversity index (richness and evenness) and Chao1 index (richness). Significant differences in the number of OTUs and alpha diversity metrics between samples were determined using paired t-tests. Beta diversity was explored by non-metric multidimensional scaling (nMDS) of Bray-Curtis dissimilarity among samples. Statistical significance was calculated by permutational analysis of variance (PERMANOVA). Venn diagrams were visualized with the R package ‘VennDiagram’ (Chen and Boutros 2011), illustrating the shared and unique genera among each bathroom paired sample. Linear discriminant analysis (LDA) effect size (LEfSe) was applied to identify specific bacterial genera between the samples (Segata *et al*. 2011). Taxa were considered significant based on LDA scores greater than 4.0 and p-values smaller than 0.05.

### Possibility of pathogens in the bathroom

The presence or absence of genera on the National Institute of Infectious Diseases (NIID) in Japan list of Bio Safety Level (BSL) 2 and 3 were checked in each sample (for the genus in NIID list, see Supplemental Table S4, and for the list of species, see the following link: https://www.nite.go.jp/nbrc/mrinda/list/risk/bacteria/ALL, updated 2010-06). As a result, each read corresponding to BSL2 and BSL3 species was collected, and then sequences longer than 1,000 bp were filtered by seqkit (v0.8.0). The sequences were clustered with psi-cd-hit.pl to identify representative sequences (Li and Godzik 2006; Fu *et al*. 2012). The settings in psi-cd-hit.pl were the default except for -prog blastn. For further analysis, the sequences with the highest number of reads in each cluster were used. Taxonomy was identified with BLASTn (E value = 1.0e-10, word size = 7, reward = 2, penalty = −3, gap open = 5, gap extend = 2, filter: unmarked “low complexity regions”) (Camacho *et al*. 2009) against 16S ribosomal RNA from curated type strain sequences from bacteria and archaea in GenBank (update date: 2021/01/11, number of sequences: 21,699). The analysis scheme is shown in Figure 1C.

## Supporting information

SupTableS4

SupFigS1

SupTableS1

SupTableS2

SupTableS3

## Data availability

The authors declare that all the data supporting the findings of this study are available within the paper (and its Supplemental files) and that raw data were presented where possible. The raw MiSeq data reported in the paper (Figure 6, Supplemental Figures and Tables) have been uploaded to the DDBJ database under the accession number DRA010182. It will be open to public when this manuscript is accepted for publication

## Author contributions

**S.F.** and **F.M.** conceived and designed the experiment. **S.F.** and **F.M.** collected samples. **S.F.** performed the experiments and analysed the data. **S.F.**, **Y.N.**, and **F.M.** interpreted the data. **S.F.** wrote the manuscript. **S.F.**, **Y.N.**, and **F.M.** reviewed drafts of the manuscript.

## Acknowledgements

This work was supported by The Kyoto University Research Funds for Young Scientists and the Japan Society for the Promotion of Science under Grants-in-Aid for Scientific Research (KAKENHI) (grant number 20K18903 to S.F.). KAKENHI (grant numbers 18K19674/18KK0436/20H00562) and the Japan Agency for Medical Research and Development (grant number 20wm0225012h0001/21fk0108129h0502) awarded grants to F.M. This manuscript was edited for English language by American Journal Experts (AJE).

---

**Supplemental Figure S1**. Taxonomic composition of the bacterial community in each sample at the genus level using a stacked bar plot (top 30 genera).

