## Supplementary figures and images for "Structure of bacterial communities in Japanese-style bathrooms: Comparative sequencing of bacteria in shower water and showerhead biofilms using a portable nanopore sequencer"

### SupFigS1

#### Slide 1
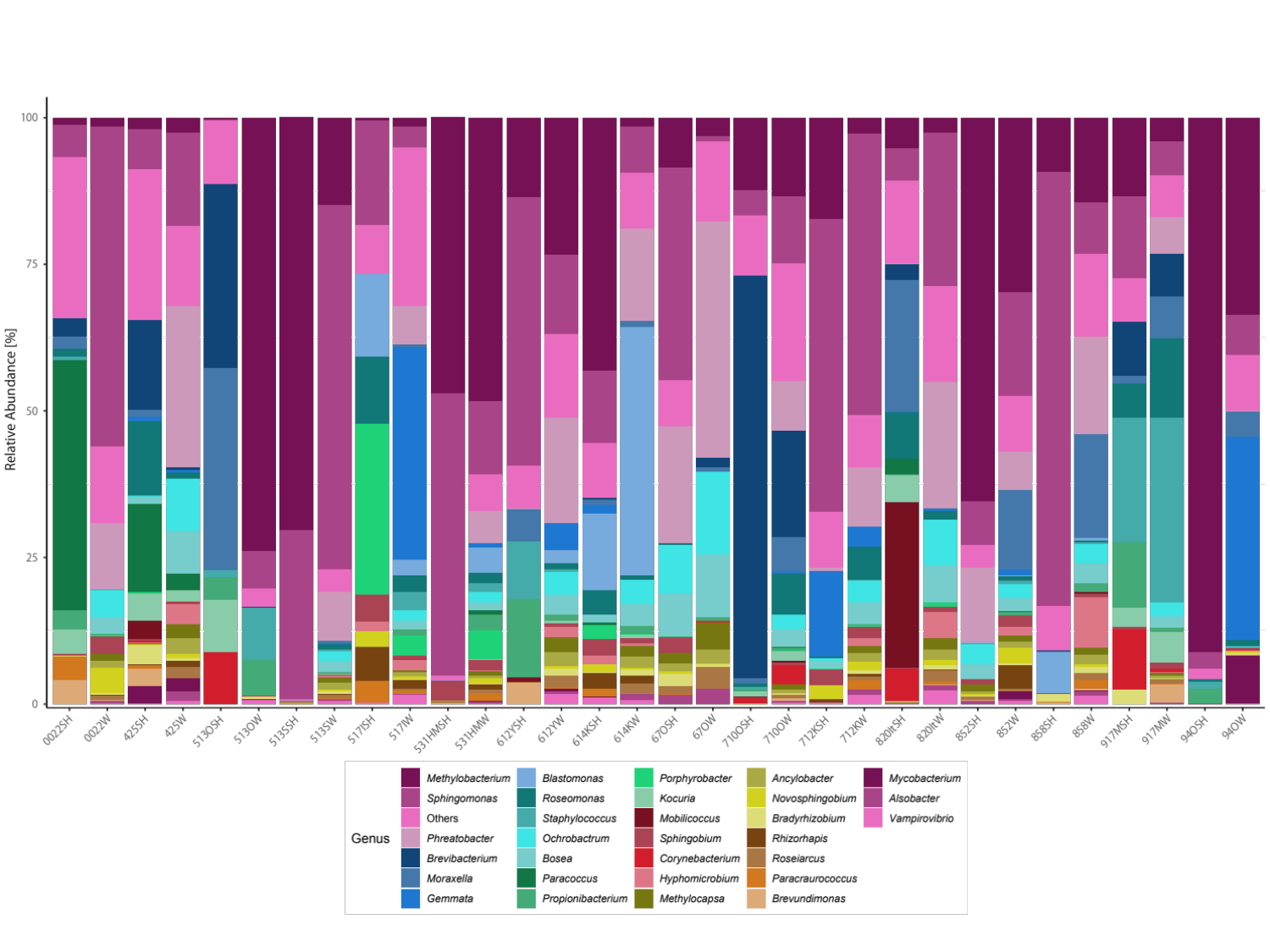
